## Supplementary material for "Cryo-EM structures of the *E. coli* Ton and Tol motor complexes": Tables

### Table 1: CryoEM data collection, structure determination and model statistics.

| | **TonB-ExbBD** | | | **$TolA_{TEV}QR$** | | **TolAQR** |
|---|---|---|---|---|---|---|
| ***Data collection*** | | | | | | |
| Nominal magnification | 105,000x | | | 105,000x | | 45,000x |
| Voltage (kV) | 300 | | | 300 | | 200 |
| Exposure time (s/frame) | 0.05 | | | 0.075 | | 0.08 |
| Number of frames | 46 | | | 30 | | 30 |
| Total dose (ē/Å$^2$) | 69.9 | | | 61.4 | | 69.9 |
| Defocus range (μm) | -0.8 to -2.8 | | | -0.8 to -2.8 | | -0.7 to -2.5 |
| Pixel size (Å) | 0.83 | | | 0.83 | | 0.89 |
| ***Image processing*** | | | | | | |
| Micrographs selected | 5,784 | | | 7,811 | | 5,084 |
| Initial particle images (no.) | 5,414,663 | | | 11,112,982 | | 1,519,049 |
| | TonB bound to $ExbB_C$ | TonB bound to $ExbB_E$ | TonB bound to $ExbB_A$ | $TolA_{TEV}QR$-I | $TolA_{TEV}QR$-II | |
| Final particle images (no.) | 227,584 | 30,556 | 40,906 | 59,369 | 78,674 | 107,631 |
| Symmetry imposed | C1 | C1 | C1 | C1 | C1 | C1 |
| FSC threshold | 0.143 | 0.143 | 0.143 | 0.143 | 0.143 | 0.143 |
| Final map resolution (Å) | 2.80 | 3.16 | 3.19 | 2.94 | 3.18 | 5.40 |
| Resolution range (Å) | 2.7-3.5 | 2.9-4.5 | 3.0-4.4 | 2.7-3.9 | 2.9-4.5 | |
| ***Atomic model*** | | | | | | |
| Number of protein residues | 1209 | 1199 | 1199 | 1221 | 1206 | |
| ***Validation*** | | | | | | |
| Most favored (%) | 99.50 | 99.07 | 99.24 | 99.25 | 98.57 | |
| Allowed (%) | 0.50 | 0.93 | 0.76 | 0.75 | 1.43 | |
| Disallowed (%) | 0 | 0 | 0 | 0 | 0 | |
| Rotamer outliers (%) | 0.53 | 0.43 | 0.43 | 0.30 | 0.51 | |
| r.m.s.d Bond lengths (Å) | 0.011 | 0.011 | 0.011 | 0.012 | 0.012 | |
| r.m.s.d Bond angles (°) | 1.626 | 1.624 | 1.603 | 1.628 | 1.639 | |
| Clashscore | 0.48 | 2.40 | 1.26 | 1.91 | 2.03 | |
| Map CC (mask) | 0.88 | 0.88 | 0.88 | 0.86 | 0.84 | |
| Map CC (volume) | 0.88 | 0.88 | 0.88 | 0.85 | 0.83 | |
| ***Deposition ID*** | | | | | | |
| PDB ID | 9DDO | 9DDP | 9DDQ | 9DDM | 9DDN | |
| EMDB ID | EMD-46778 | EMD-46779 | EMD-46780 | EMD-46776 | EMD-46777 | |

**Table 2: plasmids.**

| | | |
|---|---|---|
| $ptolA_{strep}$ | *tolA* N-term streptag-II, pCDFDuet, $Sm^r$ | *this study* |
| $ptolA_{strep-TEV}$ | $ptolA_{strep}$ TEV site, pCDFDuet, $Sm^r$ | *this study* |
| pQR | *ybgC-tolQ-tolR* operon, pT7-1QR, $Amp^r$ | *Germon et al., 1998*[1] |
| $ptonB_{strepII}$ | *tonB* C-term TEV site and streptag-II, pACYCDuet, $Cm^r$ | *this study* |
| pexbB | *exbB*, pET26b, $Kan^r$ | *Celia et al., 2016*[2] |
| $pexbD_{10his}$ | *exbD*, C-term TEV site and 10his-tag, pCDFDuet, $Sm^r$ | *Celia et al., 2016*[2] |

**Table 3: Interface buried surface areas (A$^2$)**

| | TonB-ExbBD |
|---|---|
| $ExbB_{D-C}$ | 1551.2 |
| $ExbB_{B-A}$ | 1529.0 |
| $ExbB_{E-A}$ | 1485.4 |
| $ExbB_{C-B}$ | 1392.0 |
| $ExbB_{E-D}$ | 1378.4 |
| TonB-$ExbB_{C}$ | 477.6 |

| | $TolA_{TEV}QR$-I | $TolA_{TEV}QR$-II |
|---|---|---|
| $TolQ_{D-C}$ | 1041.9 | 981.5 |
| $TolQ_{B-A}$ | 1092.1 | 1053.9 |
| $TolQ_{E-A}$ | 1254.3 | 1275.3 |
| $TolQ_{C-B}$ | 1381.1 | 1368.1 |
| $TolQ_{E-D}$ | 984.1 | 907.8 |
| $TolA_{F}$-$TolQ_{C}$ | 624.8 | 472.3 |
| $TolA_{F}$-$TolQ_{B}$ | 135.5 | 110.2 |
| $TolA_{F}$-$TolQ_{CB}$ | 760.3 | 582.5 |
| $TolA_{G}$-$TolQ_{E}$ | 599.1 | 606.1 |
| $TolA_{G}$-$TolQ_{D}$ | 196.7 | 180.1 |
| $TolA_{G}$-$TolQ_{ED}$ | 795.8 | 786.2 |

Interfaces calculated with PISA[3]


1. Germon P, Clavel T, Vianney A, Portalier R, Lazzaroni JC. Mutational analysis of the Escherichia coli K-12 TolA N-terminal region and characterization of its TolQ-interacting domain by genetic suppression. *J Bacteriol* **180**, 6433-6439 (1998).

2. Celia H*, et al.* Structural insight into the role of the Ton complex in energy transduction. *Nature* **538**, 60-65 (2016).

3. Krissinel E. Stock-based detection of protein oligomeric states in jsPISA. *Nucleic Acids Res* **43**, W314-319 (2015).
