## Supplementary figures for "Cryo-EM structures of the *E. coli* Ton and Tol motor complexes"

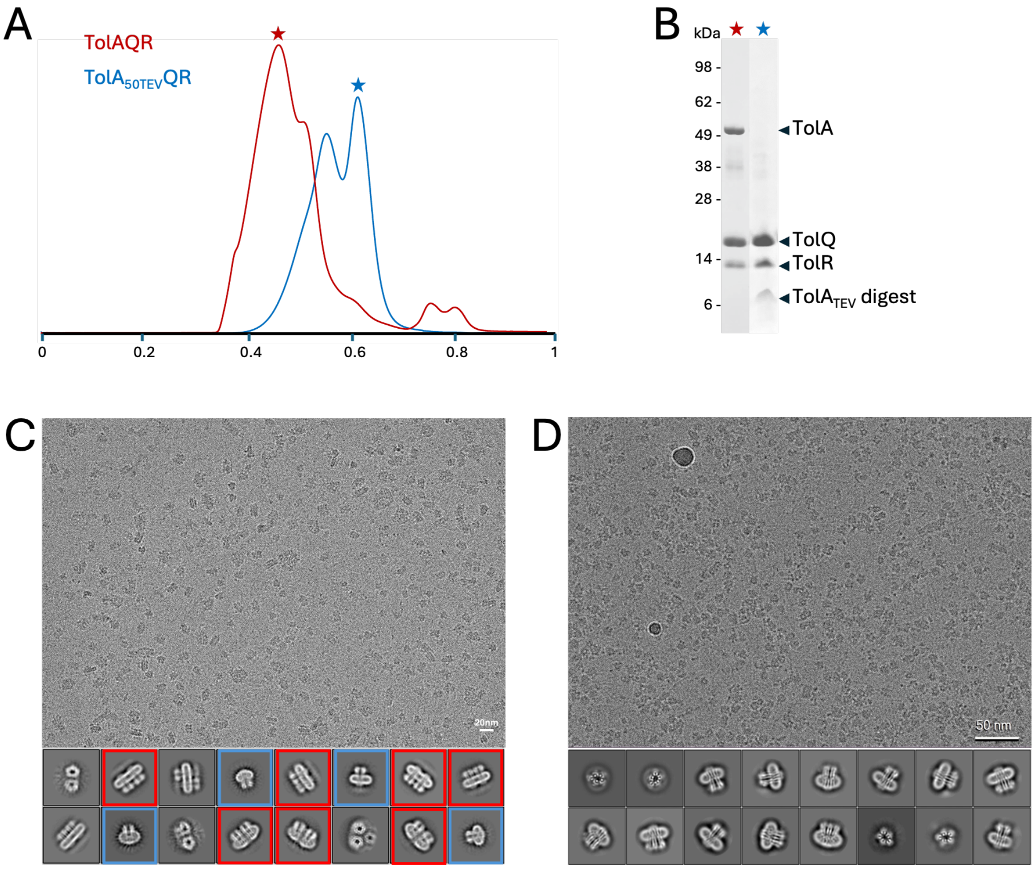


**Supplementary figure 1**

A. SEC profiles of TolAQR (red) and $TolA_{TEV}QR$ after TEV digestion (blue). The horizontal axis represents elution in column volume, the vertical axis is the relative absorbance at 280nm.
B. Coomassie stained SDS-Page of aliquots of SEC at positions indicated with stars in A. The bands corresponding to TolA, TolQ, TolR and the digest of $TolA_{TEV}$ are indicated.
C. Electron micrograph of frozen TolAQR in LMNG, showing a variety of oligomers. Lower panel: representative 2D classes of TolAQR showing monomers of TolAQR (blue), dimers (red), and higher oligomers.
D. Electron micrograph of frozen TEV digested $TolAQR_{TEV}$ in LMNG. Lower panel: representative 2D classes showing monomers of $TolAQ_{TEV}$ in different orientations.

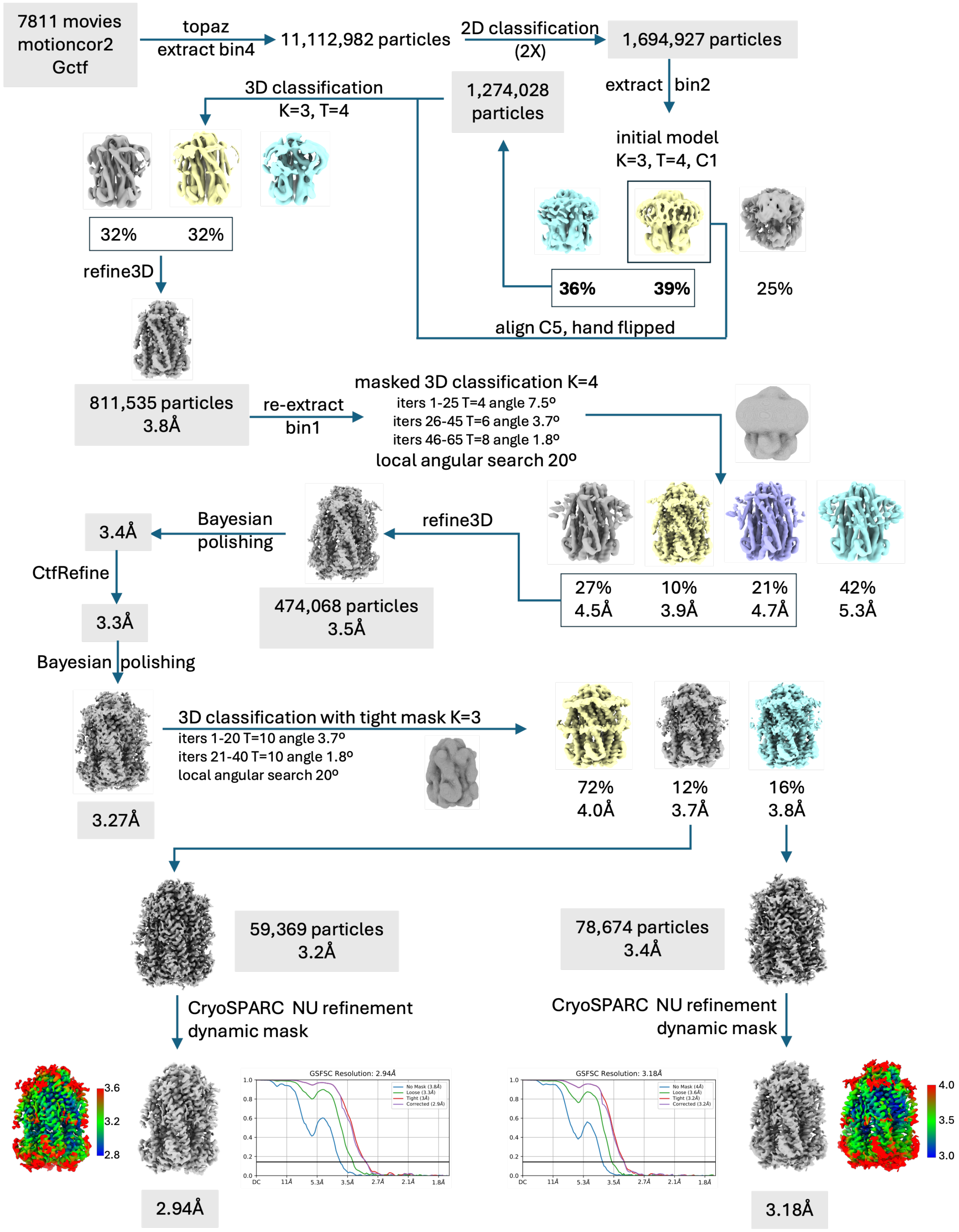


**Supplementary figure 2**

Schematic diagram of the cryoEM data processing procedures for TEV digested TolAQR$_{TEV}$.

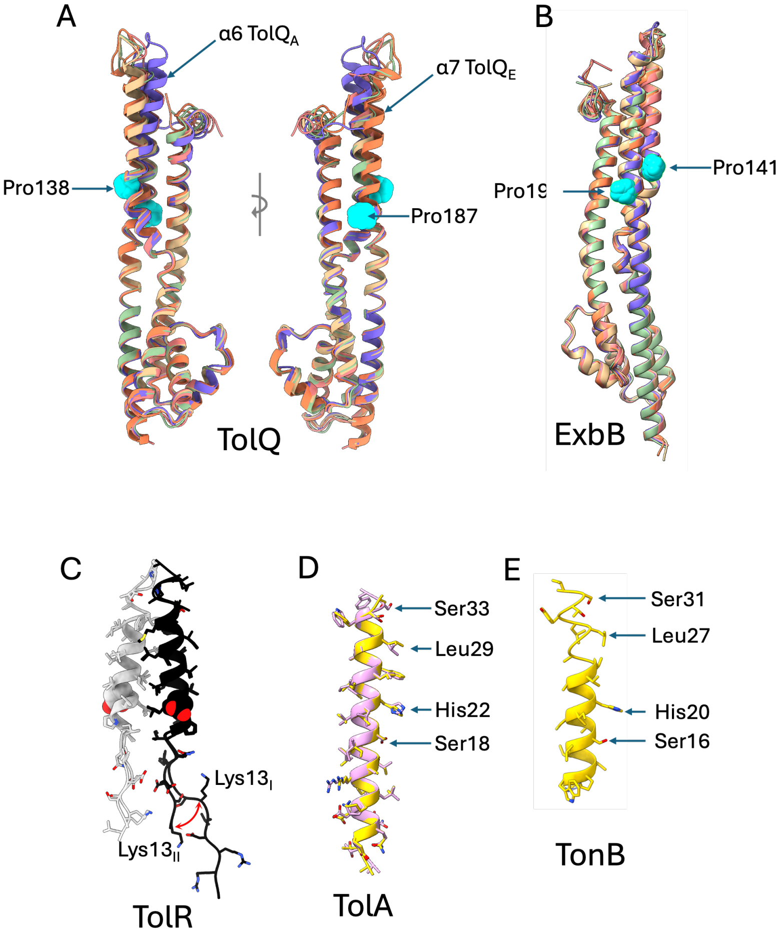


**Supplementary figure 3**

A. Superimposition of the five TolQ chains in cartoon representation and observed in two orientations. The indicated α-6 helix of $TolQ_A$ (blue) is tilted towards α-2 and α-1, while α-7 of $TolQ_E$ (orange) protrudes inside the hydrophobic pore. The conserved proline 138 and 187 that form kinks in α-6 and α-7 are represented as spheres and colored cyan.
B. Superimposition of the five ExbB chains (A to E) of the TonB-ExbBD complex, showing they share the same conformation. The conserved prolines 141 and 190 are shown as spheres and colored in cyan.
C. Superimposition of the $TolR_Y$ (black) and $TolR_Z$ (white) from the structures at 2.9Å ($TolA_{TEV}$QR-I) and 3.2Å ($TolA_{TEV}$QR II). The curved double arrow shows the swing in orientation of the residue Lys13. The essential Asp23 are shown as spheres.
D. Superimposition of $TolA_F$ (gold) and $TolA_G$ (plum). The two chains have the same conformation. The Ser18, His22, Leu29 and Ser33 that form the SHLS motif are indicated.
E. Cartoon representation of TonB. The Ser16, His20, Leu27 and Ser31 that form the SHLS motif are indicated.

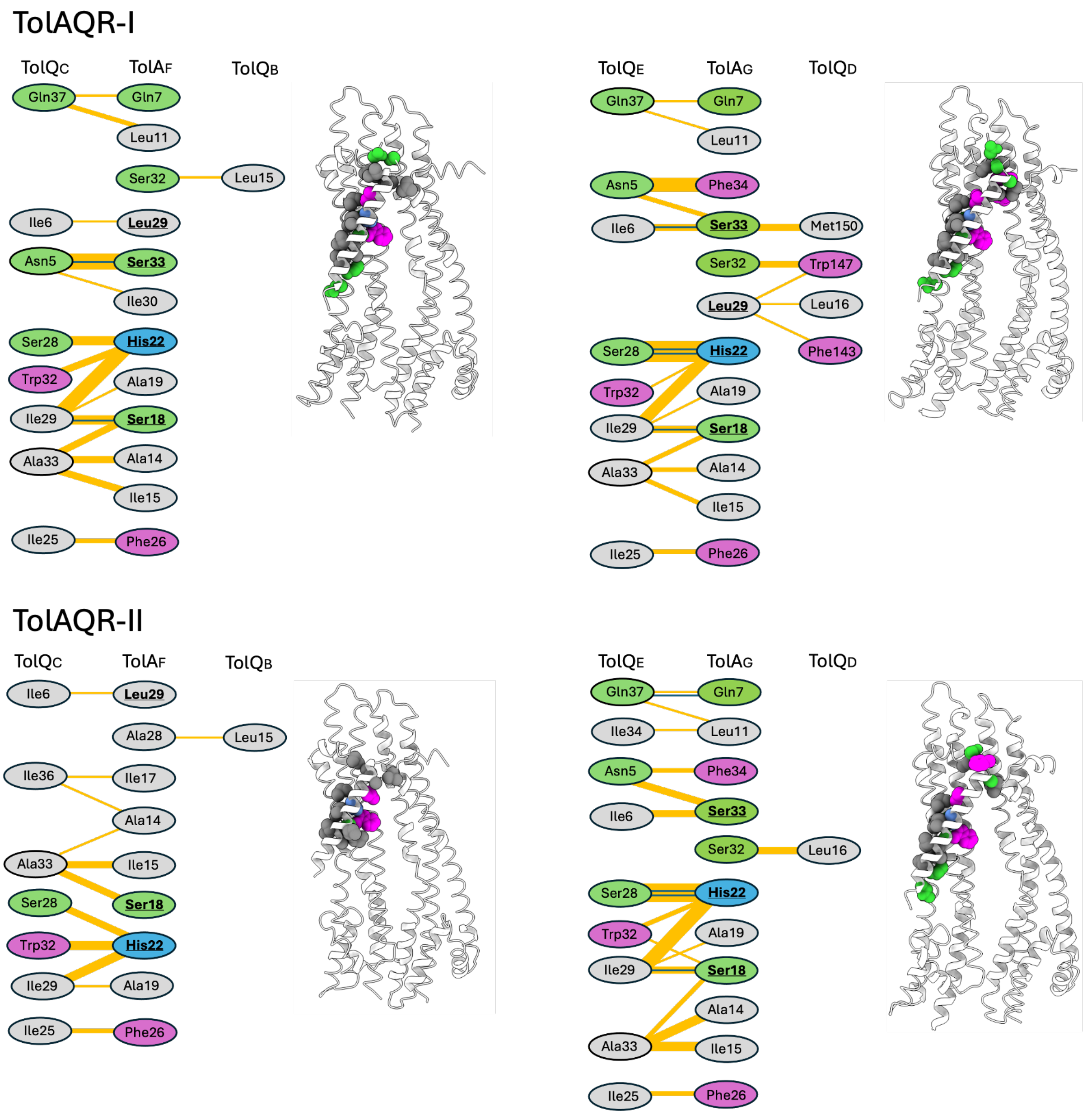
TolAQR-I
TolQC
TolAF
TolQB
Gln37
Gln7
Leu11
Ser32
Leu15
Ile6
Leu29
Asn5
Ser33
Ile30
Ser28
His22
Trp32
Ala19
Ile29
Ser18
Ala33
Ala14
Ile15
Ile25
Phe26
TolQE
TolAG
TolQD
Gln37
Gln7
Leu11
Asn5
Phe34
Ile6
Ser33
Met150
Ser32
Trp147
Leu29
Leu16
Ser28
His22
Phe143
Trp32
Ala19
Ile29
Ser18
Ala33
Ala14
Ile15
Ile25
Phe26
TolAQR-II
TolQC
TolAF
TolQB
Ile6
Leu29
Ala28
Leu15
Ile36
Ile17
Ala14
Ala33
Ile15
Ser28
Ser18
Trp32
His22
Ile29
Ala19
Ile25
Phe26
TolQE
TolAG
TolQD
Gln37
Gln7
Ile34
Leu11
Asn5
Phe34
Ile6
Ser33
Ser32
Leu16
Ser28
His22
Trp32
Ala19
Ile29
Ser18
Ala14
Ala33
Ile15
Ile25
Phe26

### TonB-ExbBD

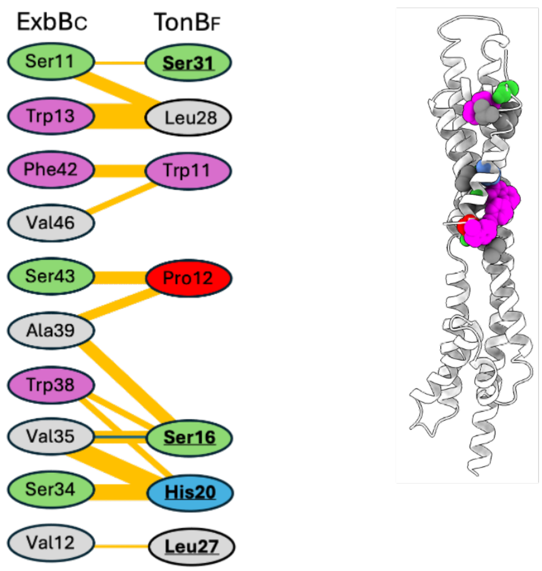


**Supplementary figure 4**

Interactions between TolA and TolQ, and TonB and ExbB as determined with PDBsum[1] (https://www.ebi.ac.uk/thornton-srv/databases/pdbsum/). For each set of interactions, a cartoon representation of the subunits listed is presented, with the side chains of the interacting residues shown as spheres and colored accordingly. The residues in bold and underlined are part of the SHLS motif. Hydrogen bonds are shown with blue lines. Non bonded contacts are shown with orange lines and the width of the line is proportional to the number of atomic contacts.

Residue colors: Positive (H, K, R); negative (D, E); neutral (S, T, N, Q), aliphatic (A, V, L, I, M), aromatic (F, Y, W)

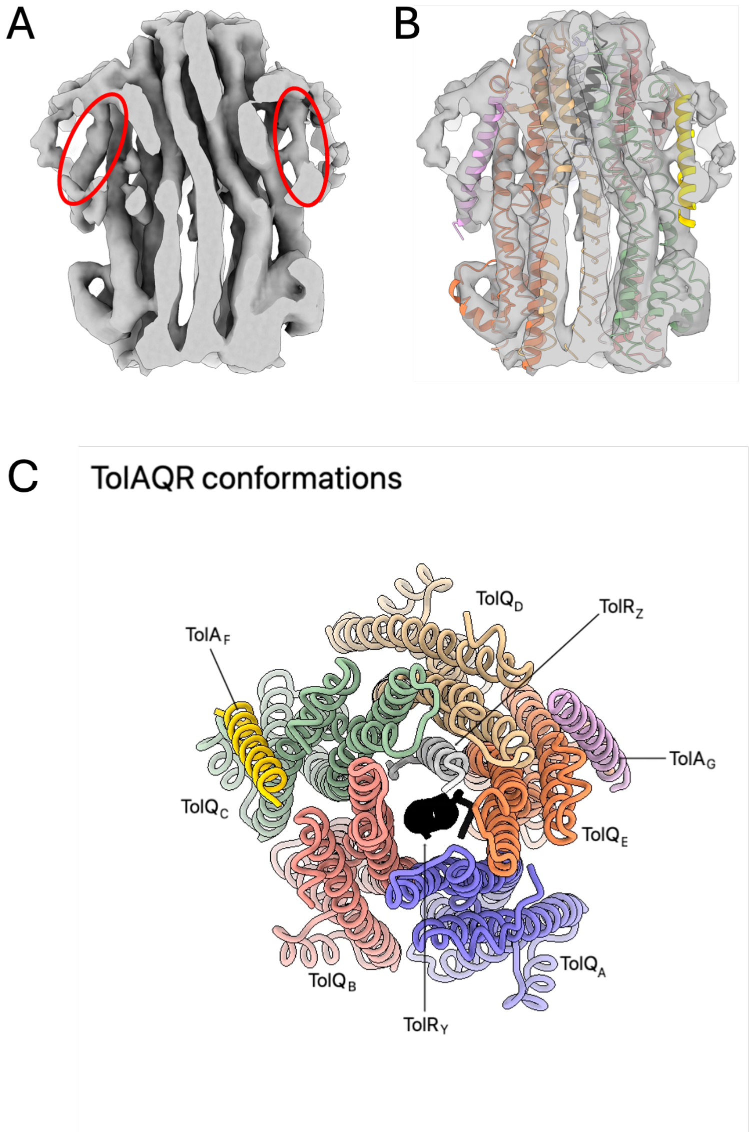


**Supplementary figure 5**

A. CryoEM 3D map represented as isosurface of the intact TolAQR in LMNG at 5.2Å resolution. The elongated densities corresponding to the TolA TMs are circled in red.

B. Same as A, the 3D map is semitransparent, and superimposed with the TolAQR model represented as cartoon, showing the match between the two structures.

C. Movie: morph between the TolA$_{TEV}$QR-I and -II structures at 2.9 and 3.2Å resolution, highlighting the conformational transitions.

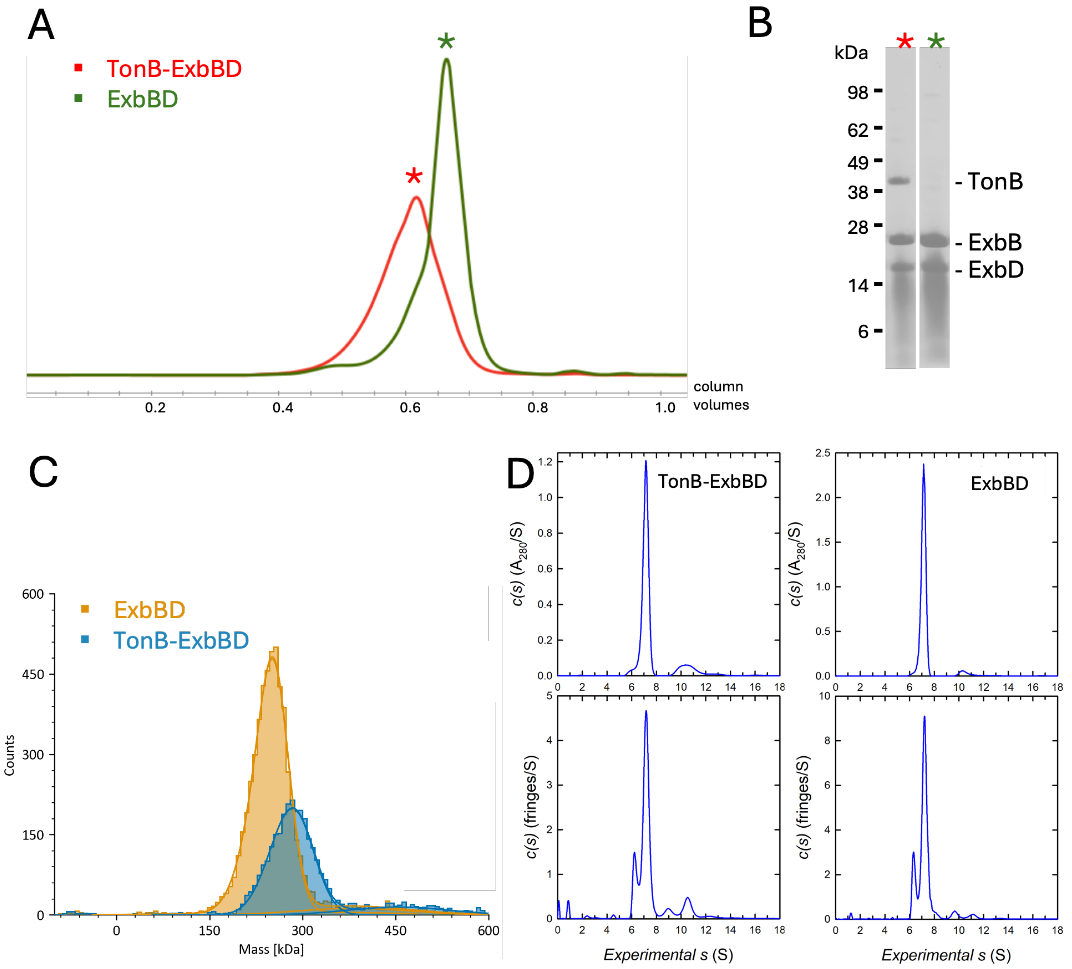


**Supplementary figure 6**

A. SEC elution profile of TonB-ExbBD (red) and ExbBD (green) in PMAL-C12. The horizontal axis corresponds to elution in column volumes, the vertical axis is the relative absorbance at 280nm.

B. Coomassie stained SDS-Page of an aliquot of purified TonB-ExbBD (left lane) and ExbBD (right lane) in PMAL-C12. The bands corresponding to TonB, ExbB and ExbD are indicated. The smearing pattern below the ExbD bands is due to the presence of PMAL-C12.

C. Mass distribution profile of ExbBD (orange) and TonB-ExbBD (blue) in PMAL-C12 as measured by mass photometry. The horizontal axis represents the mass in kilodalton (kDa), the vertical axis the count of species hitting the surface of the coverslip. The ExbBD peak is centered at 248 kDa, TonB-ExbBD at 282 kDa.

D. Sedimentation velocity absorbance (top panels) and interference (bottom panels) profiles for TonB-ExbBD (left panels) and ExbBD (right panels) in PMAL-C12 showing a major species at 7.1 S corresponding to the expected complexes.

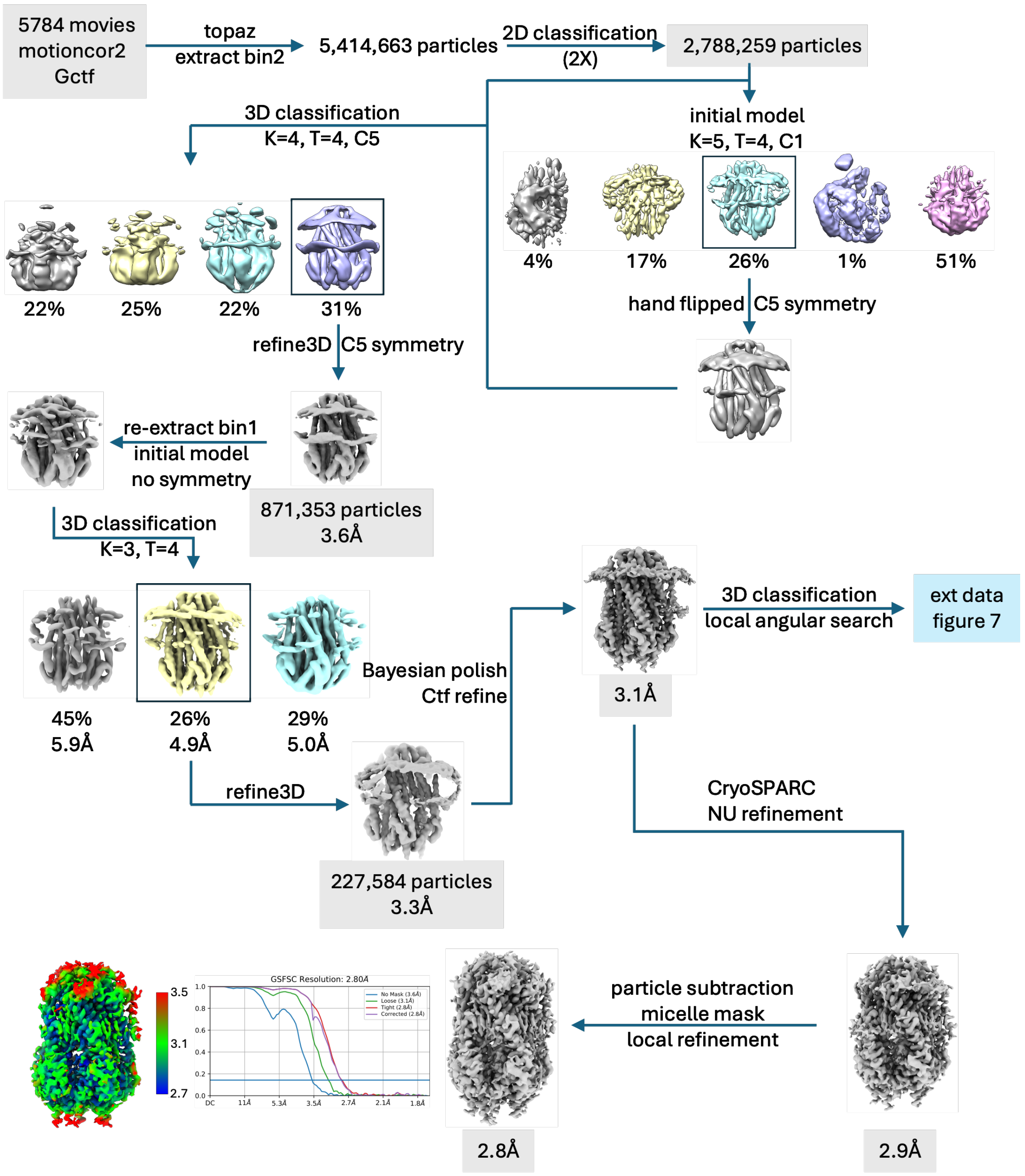


**Supplementary figure 7**

Schematic diagram of the cryoEM data processing procedures for TonB-ExbBD resulting in the consensus map at 2.8Å resolution.

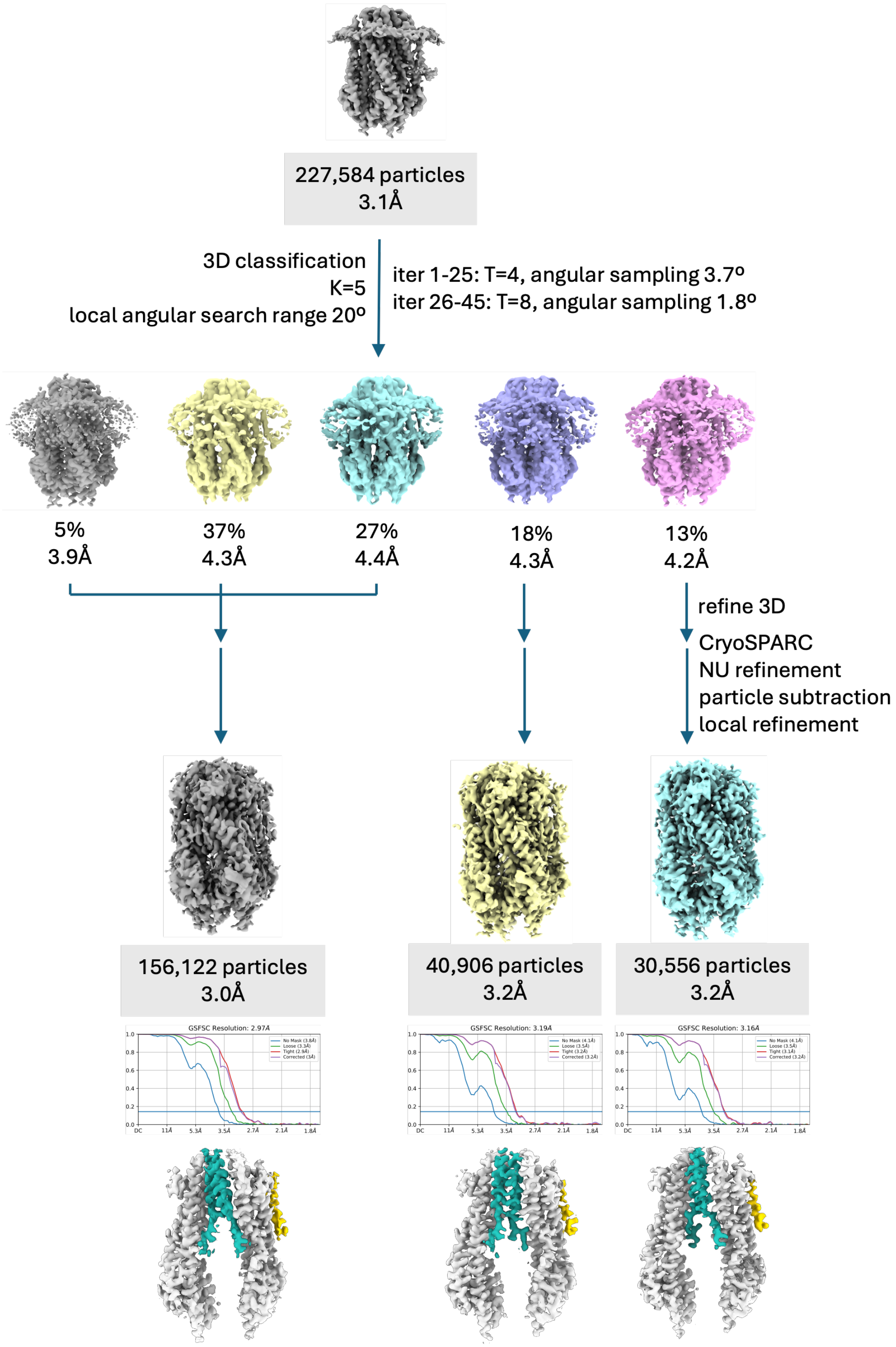


**Supplementary figure 8**

Schematic diagram of the cryoEM data processing procedure leading to the three structures with TonB binding to different ExbB chains.

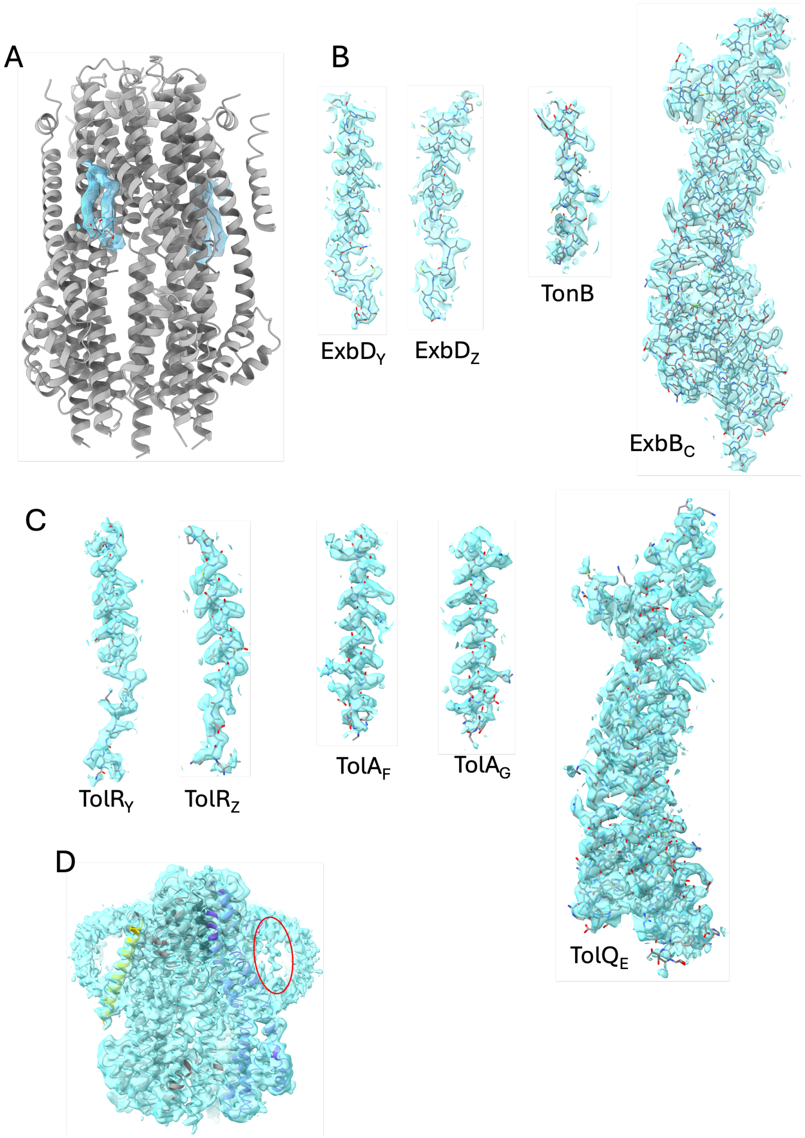


**Supplementary figure 9**

Fit between the structures and their experimentals maps. Structures of different chains are shown as ball and stick and the corresponding 3D density maps are shown as semitransparent blue isosurfaces.

A. Phospholipids: TonB-ExbBD is represented as cartoon, the two modeled phosphatidyl ethanolamine molecules are shown

B. Fit between the structures of ExbD$_Y$, ExbD$_Z$, TonB and ExbB$_C$ and their associated densities.

C. Fit between the structures of TolR$_Y$, TolR$_Z$, TolA$_F$, TolA$_G$ and TolQ$_E$ and their associated densities.

D. 3D densities of the unsharpened 3D map of TolA$_{TEV}$QR, superimposed with the TolAQR structure in cartoon. The red circle shows weak densities that likely correspond to an additional TolA bound to TolQ$_A$ (blue). The poor densities in this region are likely due to low occupancy.

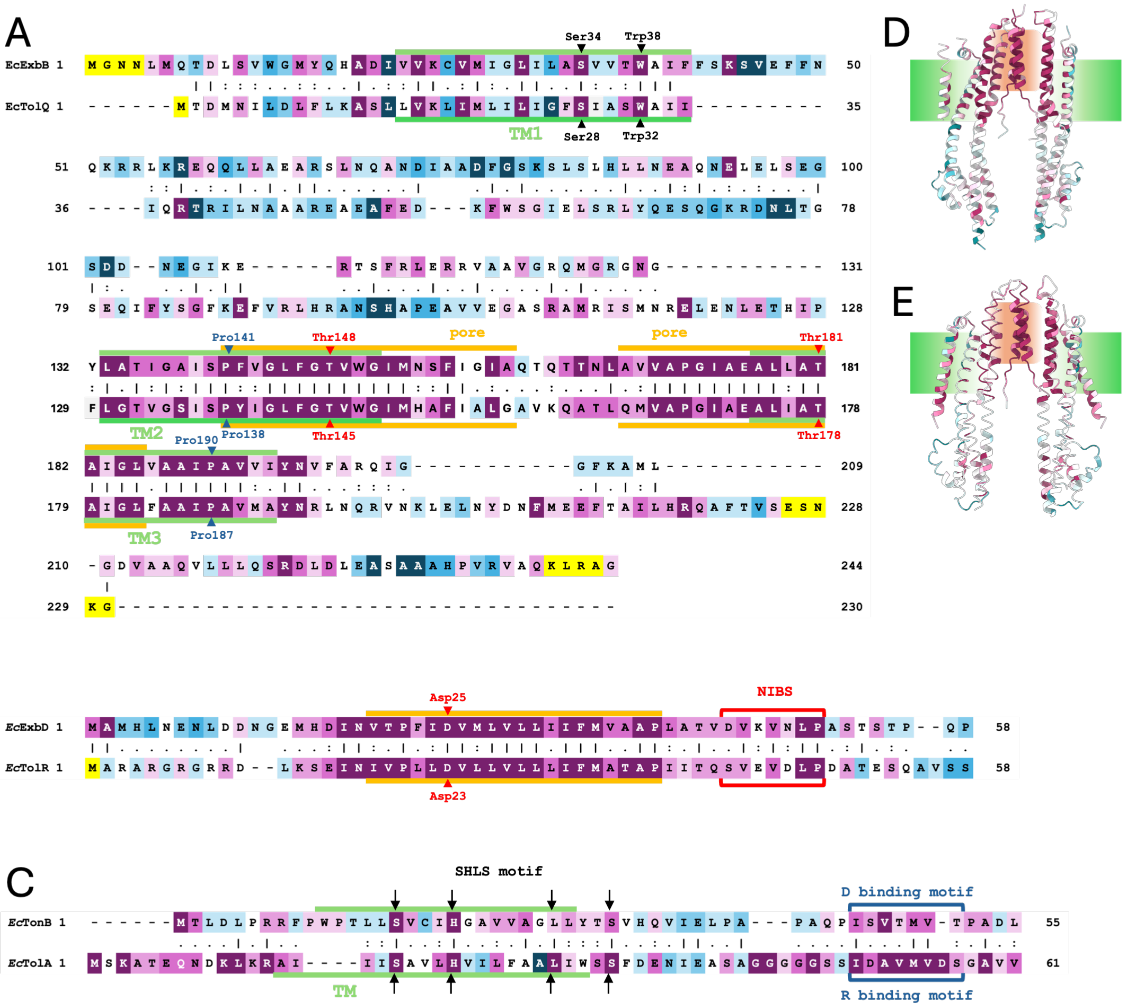


| | TonB/TolA | ExbB/TolQ | ExbD/TolR |
|---|---|---|---|
| identity | 9.8% | 27.6% | 26.1% |
| similarity | 19.2% | 39.5% | 43.8% |

**Supplementary figure 10**

Pairwise alignments[2] of *Ec*TolQ and *Ec*ExbB (A), *Ec*TolR and *Ec*ExbD (B) and *Ec*TolA and *Ec*TonB (C). “|” represent Identical residues, “:” similar residues, “.” are variable. The residues are colored according to their degree of conservation, with highly conserved residues in maroon, average in white, and poorly conserved in turquoise. The underrepresented residues are in yellow. The TM domains are shown in green, the hydrophobic pore in orange. Some conserved residues are indicated with arrowheads. For TolR/ExbD (B) and TolA/TonB (C), only the N-terminal, TM region and conserved NIBS and D-box/R-box motifs are shown. The essential Asp on ExbD and TolR (B) are indicated with arrowheads. The NIBS regions are shown with brackets. The TM domain is shown in orange as it seats in the hydrophobic pore.

The SHLS motifs on TonB and TolA (C) are shown with arrows. The D-binding motif on TonB and putative R-binding motif on TolA are shown with brackets.
D and E. Cartoon representation of TonB-ExbBD (D) and TolAQR (E) colored according to ConSurf[3]. The TolQ and ExbB chains A, B and D are omitted to show the interior of the pore. The membrane embedded regions are shown in green, the hydrophobic pore region in orange.
The table shows the identity and similarity scores for the different chains.

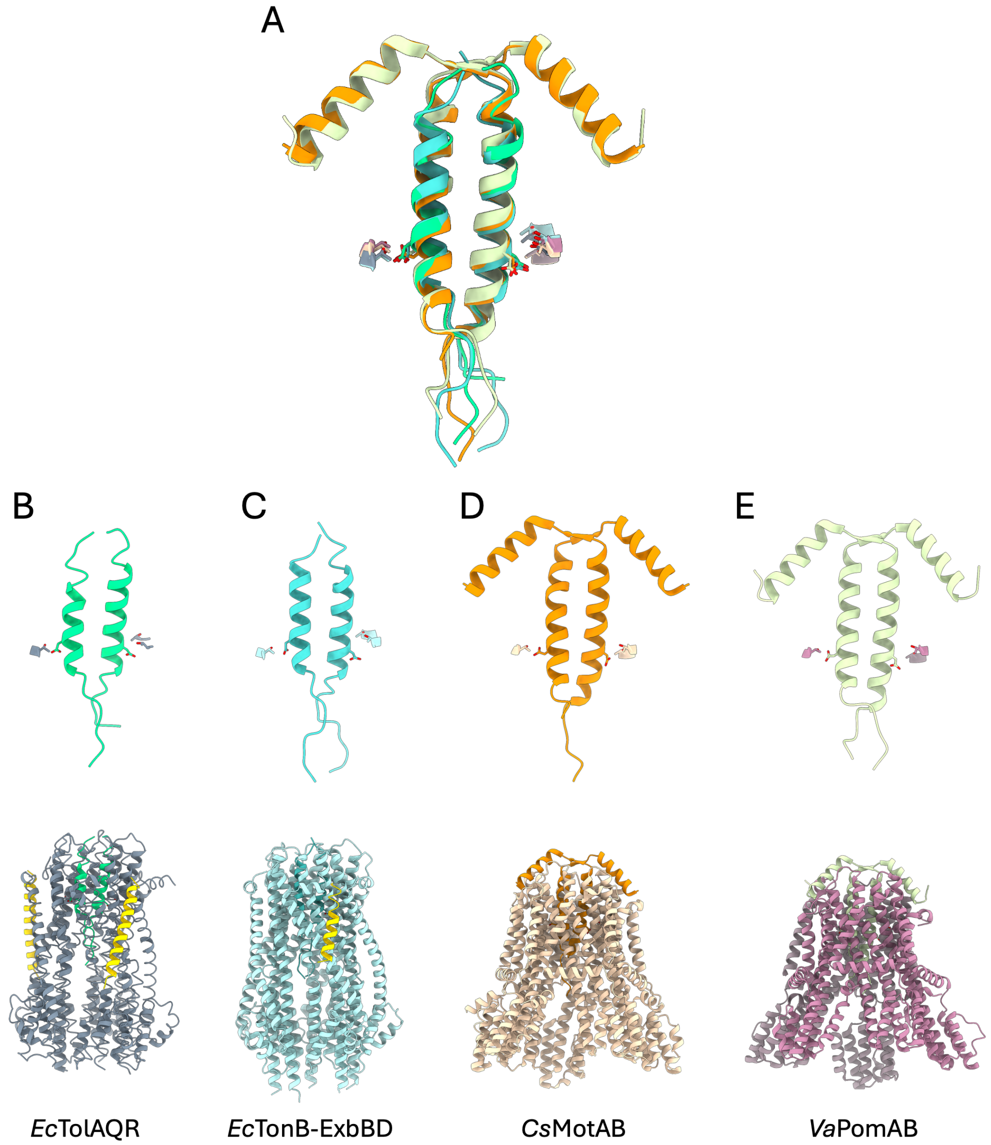


**Supplementary figure 11**

Comparison of the *Ec*TolAQR, *Ec*TonB-ExbBD, *Cs*MotAB (pdb 8UCS[4]) and *Va*PomAB structures (pdb 8BRD[5]).

A. Superimposition of cartoon representations of the TolR (green), ExbD (cyan), MotB (orange) and PomB (yellow-green) structures. The essential Asp on the TMs are shown in sticks, the neighboring conserved threonines on TM3 of ExbB and TolQ, and TM4 of MotA and PomA are shown with sticks and cartoon.

B, C, D and E: structures of *Ec*TolAQR (B), *Ec*TonB-ExbBD (C), *Cs*MotAB (D) and *Va*PomAB (E) Top panels: same view as A. Lower panel: same orientation than top panel, with all the chains of the respective structures (except for *Cs*MotAB where the FliG associated chains are omitted).

The TolA (B) and TonB (C) chains are colored gold. TolQ (B) are colored dark grey, ExbB (C) light blue, MotA (D) tan and PomA (E) plum.

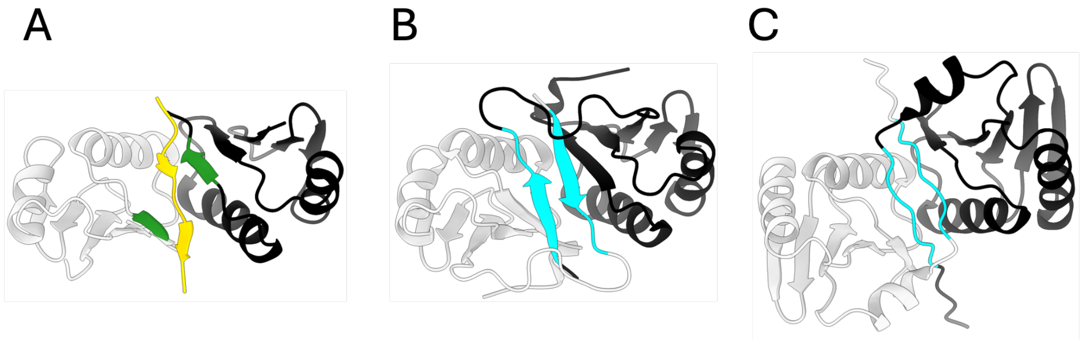


**Supplementary figure 12**

Structures of TolR and ExbD periplasmic domain dimers

A. Cartoon representation of *Ec*ExbD periplasmic dimer (black and light grey) in complex with TonB D-box peptide (gold) (pdb 8P9R[6]). The β5 strands making contacts with the D-box peptide are colored green.

B. Cartoon representation of *Sm*ExbD periplasmic dimer in the closed state (pdb 8PEK[6]). The NIBS domains are colored cyan.

C. Cartoon representation of *Ec*TolR periplasmic dimer (pdb 5BY4[7]). The NIBS domains are colored cyan.

The three dimers are viewed along their 2-fold symmetry axis.
